## Supplementary Information for "Targeted rRNA depletion enables efficient mRNA sequencing in diverse bacterial species and complex co-cultures"

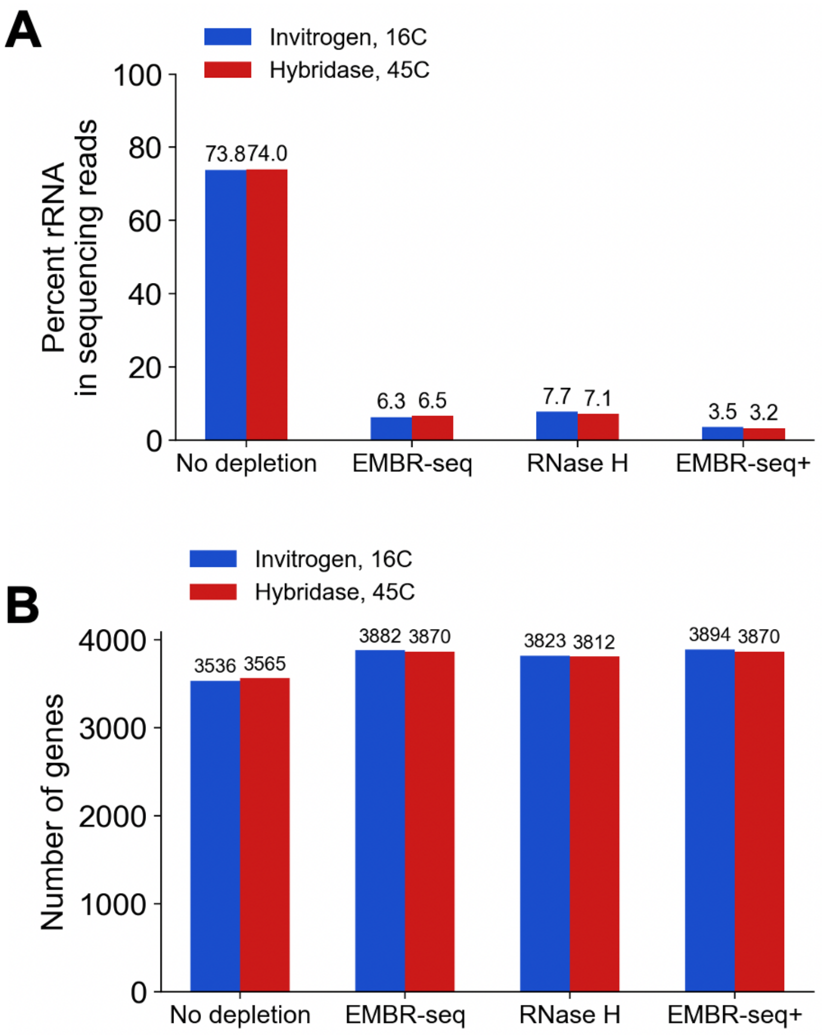


**Supplementary Figure S1. Optimization of RNase H mediated degradation of rRNA.** (**A**) Percent of sequencing reads mapping to rRNA in *E. coli* for four different library preparation methods using two different RNase H enzymes at different reaction temperatures (Invitrogen RNase H at 16°C and Hybridase Thermostable RNase H at 45°C). (**B**) The number of *E. coli* genes detected in each library. Both Invitrogen RNase H and Hybridase Thermostable RNase H show similar performance in these experiments.

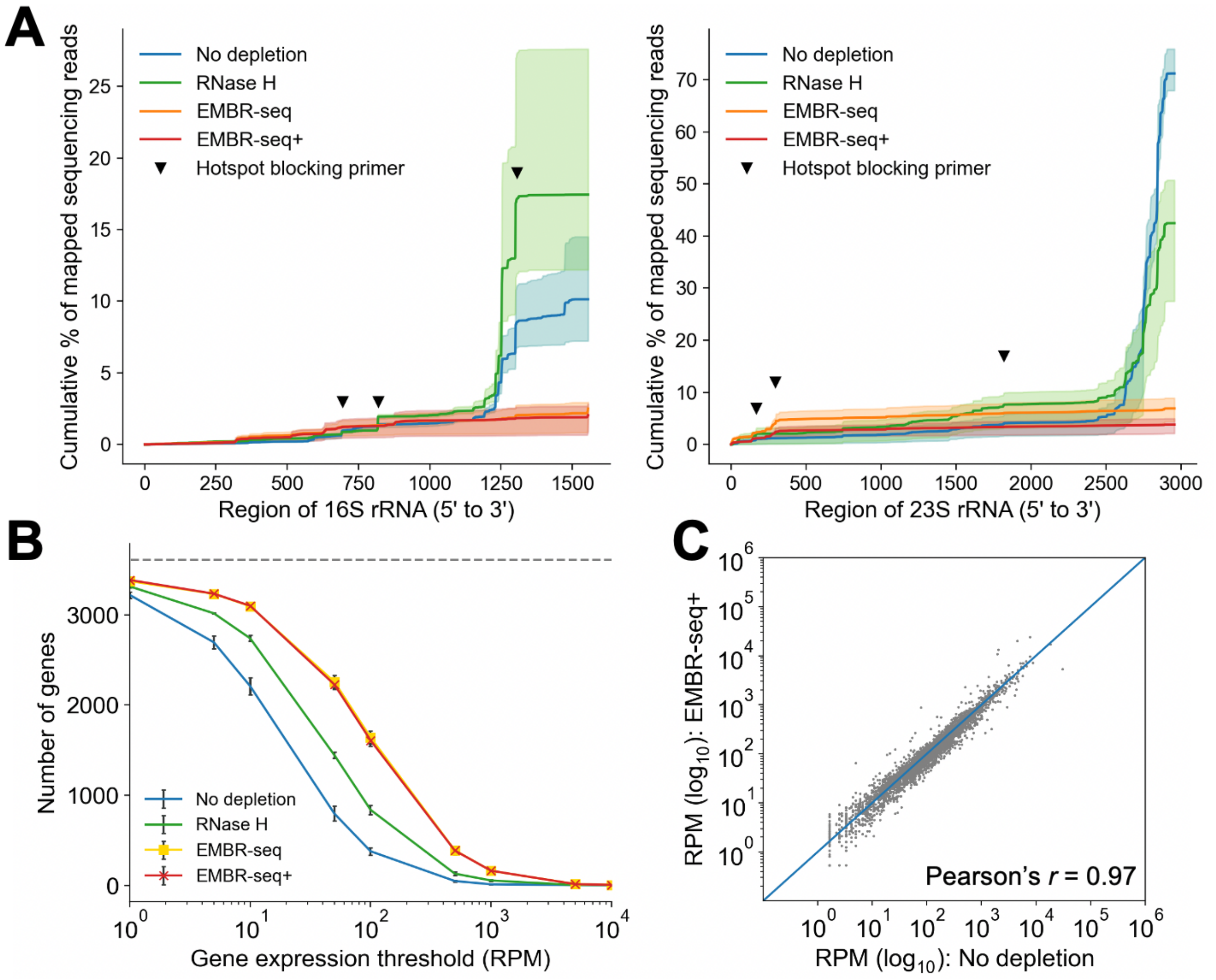


**Supplementary Figure S2. Depletion of rRNA from *G. metallireducens* using EMBR-seq+*.*** (**A**) Cumulative percentage of reads mapping along 16S and 23S rRNA of *G. metallireducens* for different depletion methods. Bold line indicates mean values and shaded regions indicate the minimum and maximum over three independent experiments. Inverted triangles indicate location of hotspots targeted by blocking primers. (**B**) Number of genes detected above different gene expression thresholds for *G. metallireducens*. Points indicate mean values and error bars indicate standard deviations over three independent experiments. (**C**) Correlation of gene expression between the "No depletion" and "EMBR-seq+" conditions for *G. metallireducens* (Pearson $r$ = 0.97). RPM is computed after removal of rRNA reads. The x- and y-coordinates of each point indicate mean values over three independent experiments in the two conditions. For consistent comparison across methods, panels a and b show data that has been downsampled to 1 million sequencing reads.

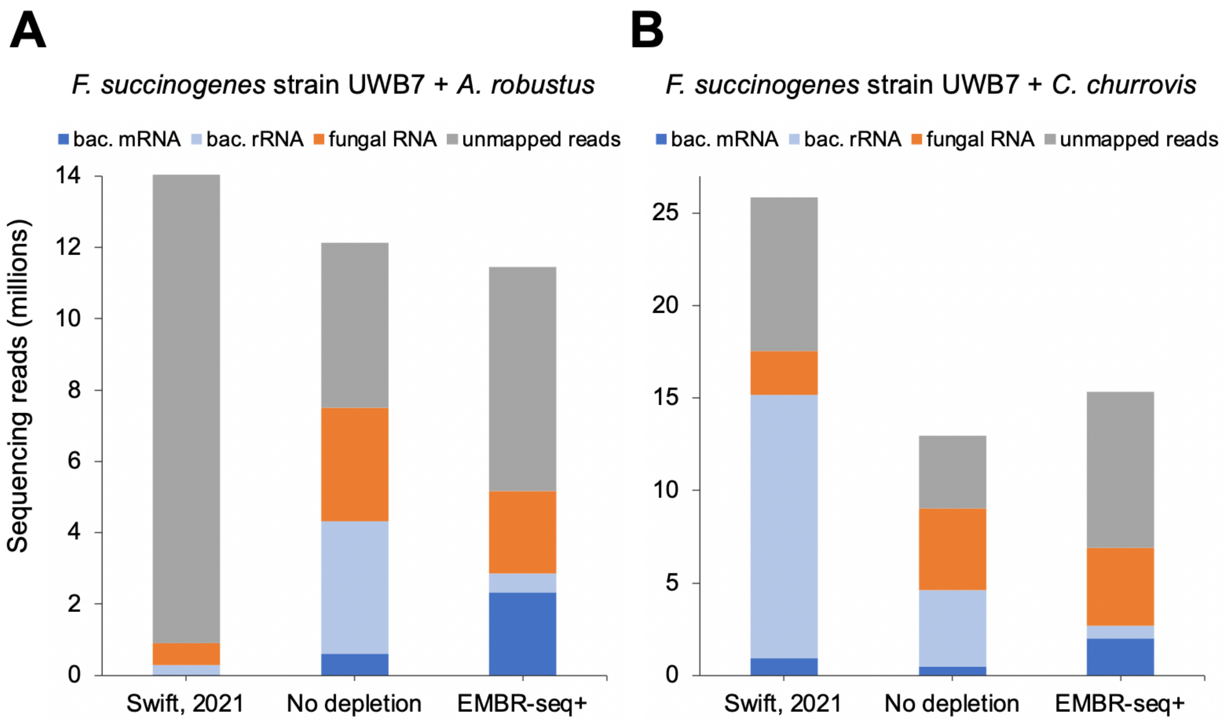


**Supplementary Figure S3. Comparison of EMBR-seq+ sequencing data to a previous study using the same co-culture systems**. "Swift, 2021" refers to data from a previous study that utilized the Ribo-Zero Gold rRNA Removal Kit (Epidemiology) for rRNA depletion. "No depletion" refers to libraries starting with "Poly-A depleted" RNA and prepared without EMBR-seq blocking primers or RNase H depletion. "EMBR-seq+" refers to "Poly-A depleted & RNase H treated" libraries. (**A**) RNA-seq libraries from co-cultures of *F. succinogenes* strain UWB7 with *A. robustus* grown on switchgrass. (**B**) RNA-seq libraries from co-cultures of *F. succinogenes* strain UWB7 with *C. churrovis* grown on switchgrass. All bars show mean values of sequencing reads over 3 or 4 replicates. To regenerate the data shown in "Swift, 2021", raw FASTQ files were downloaded from the previous study (1) and mapped to the same reference transcriptomes used throughout this study (see *RNA sequencing and bioinformatics analysis* in *Materials and Methods*).

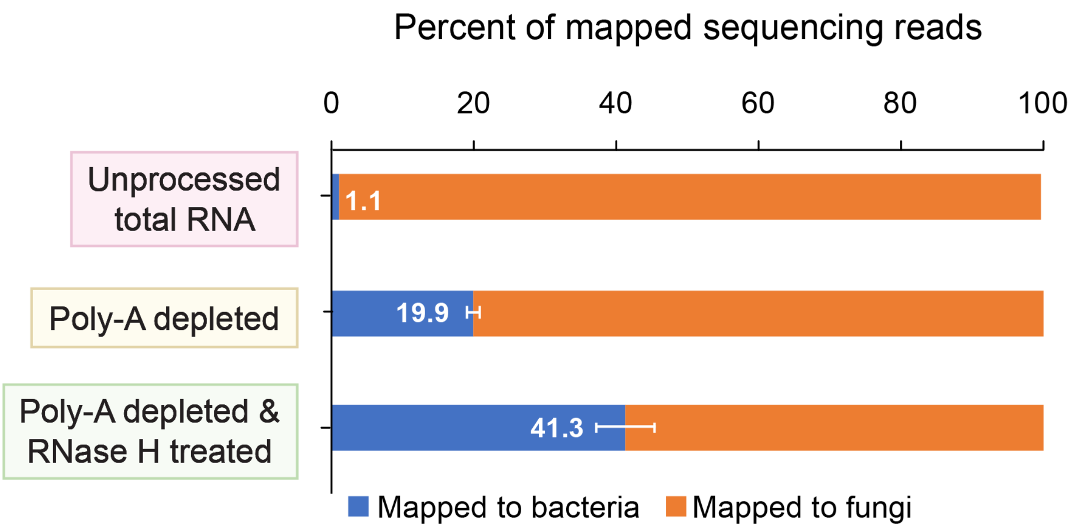


**Supplementary Figure S4. Mapped RNA reads in libraires from co-cultures between *F. succinogenes* strain UWB7 and *C. churrovis*.** Percentage of reads mapping to the bacterial and fungal genomes using the three strategies described in Figure 3A: total RNA isolated from co-culture pellets treated with EMBR-seq to remove bacterial rRNA ("Unprocessed total RNA"), fungal poly-adenylated mRNA depleted total RNA is treated with EMBR-seq to remove bacterial rRNA ("Poly-A depleted"), and "Poly-A depleted" EMBR-seq library additionally treated with RNase H to remove fungal and bacterial rRNA ("Poly-A depleted & RNase H treated"). For the "Unprocessed total RNA" library, $n$ = 1. For "Poly-A depleted" and "Poly-A depleted & RNase H treated" libraries, $n$ = 3. Bars indicate mean values and error bars indicate standard deviation from the mean.

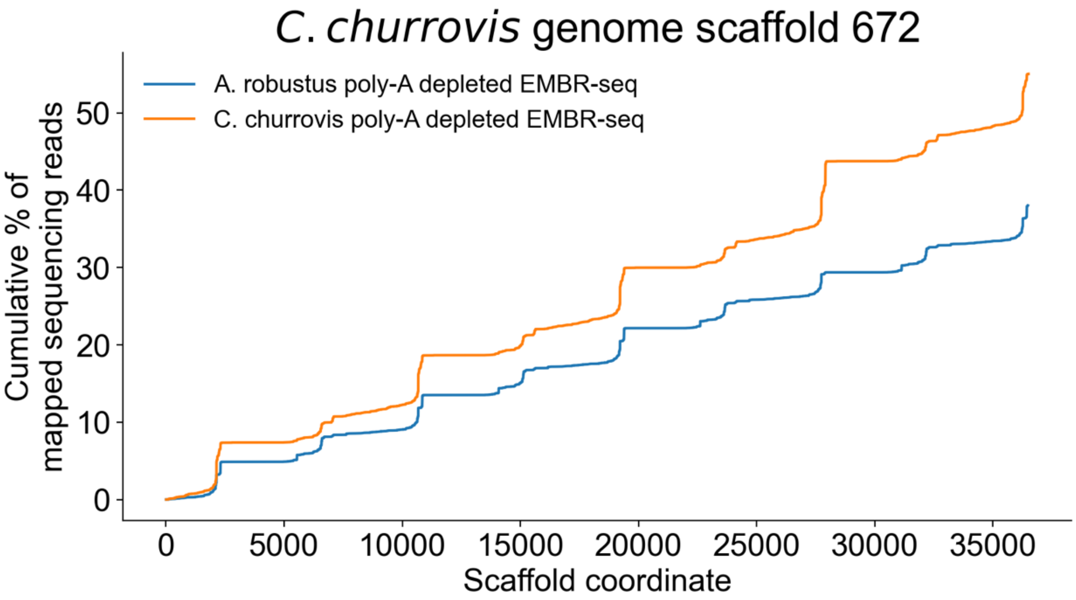


**Supplementary Figure S5. Mapping locations of sequencing reads from "Poly-A depleted" EMBR-seq libraries along *C. churrovis* scaffold 672.** Cumulative percentage of reads mapping to *C. churrovis* scaffold 672 from co-culturing *F. succinogenes* strain UWB7 with *C. churrovis* or *A. robustus* ($n$ = 1). By permitting non-unique mapping, a repetitive pattern was observed along the scaffold. EMBR-seq data from the co-culture of *F. succinogenes* strain UWB7 with *A. robustus* also exhibited high mappability on the *C. churrovis* genome, implying high sequence conservation between the two fungal strains on this scaffold.

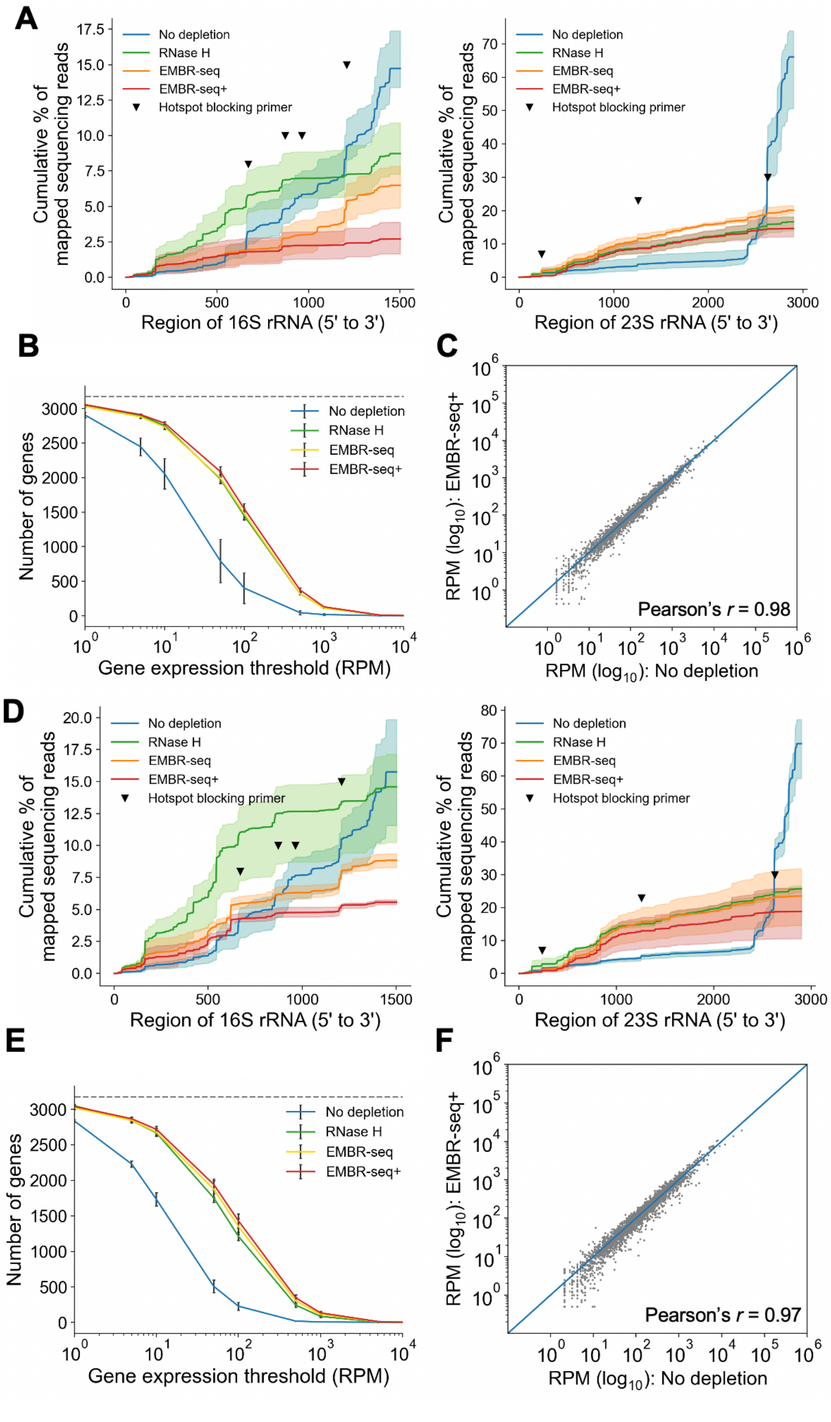


**Supplementary Figure S6. EMBR-seq+ efficiently depletes bacterial rRNA from co-cultures of *F. succinogenes* strain UWB7 with *A. robustus* or *C. churrovis*.** (**A,D**) Cumulative percentage of reads mapping to 16S and 23S rRNA of *F. succinogenes* strain UWB7 cultured with *A. robustus* (panel A) or *C. churrovis* (panel D) for different depletion methods. Bold lines indicate mean values and shaded regions indicate the minimum and maximum of three independent experiments. Inverted triangles indicate location of hotspots targeted by blocking primers. (**B,E**) Number of genes detected above different gene expression thresholds for *F. succinogenes* strain UWB7 cultured with *A. robustus* (panel B) or *C. churrovis* (panel E). Points indicate mean values and error bars indicate standard deviations over three independent experiments. (**C**,**F**) Correlation of gene expression between the "No depletion" and "EMBR-seq+" conditions for *F. succinogenes* strain UWB7 (Pearson's $r$ = 0.98 for co-culture with *A. robustus* (panel C) and $r$ = 0.97 for co-culture with *C. churrovis* (panel F)). RPM is computed after removal of rRNA reads. The x- and y-coordinates of each point indicate mean values over three independent experiments in the two conditions. For consistent comparison across methods, panels A, B, D, and E show data that has been downsampled to 1 million sequencing reads.

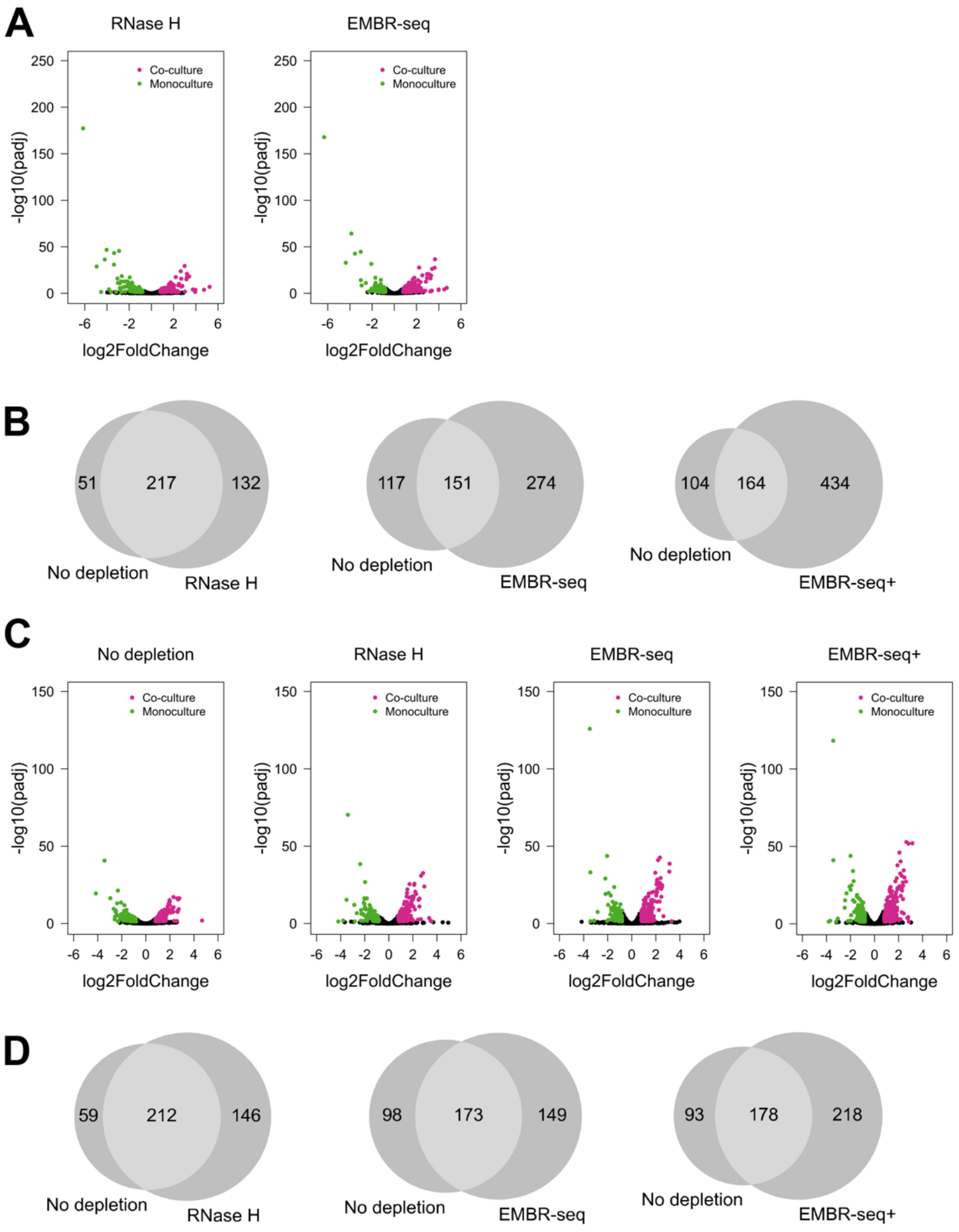


**Supplementary Figure S7. Summary of differentially expressed genes for *F. succinogenes* strain UWB7 grown in monoculture *vs.* co-culture across different depletion methods.** (**A,C**) Volcano plots of differentially expressed genes for *F. succinogenes* strain UWB7 grown in monoculture *vs.* co-culture (with *A. robustus* (panel A) or *C. churrovis* (panel C)), for the "No depletion" condition and other rRNA depletion methods. In these plots, $n = 3$ for both monoculture and co-culture conditions and all depletion methods. Colored data points indicate differentially expressed genes, determined by thresholds of $|\log_2(Fold\ Change)| > 0.8$, $P_{adj} < 0.05$, and RPM

$> 2$. Green and pink data points correspond to genes upregulated in monoculture and co-culture, respectively. (**B**,**D**) Venn diagrams show the number of differentially expressed genes identified between monoculture and co-culture (with *A. robustus* (panel B) or *C. churrovis* (panel D)) conditions that overlap between the “No depletion” condition and different rRNA depletion methods.

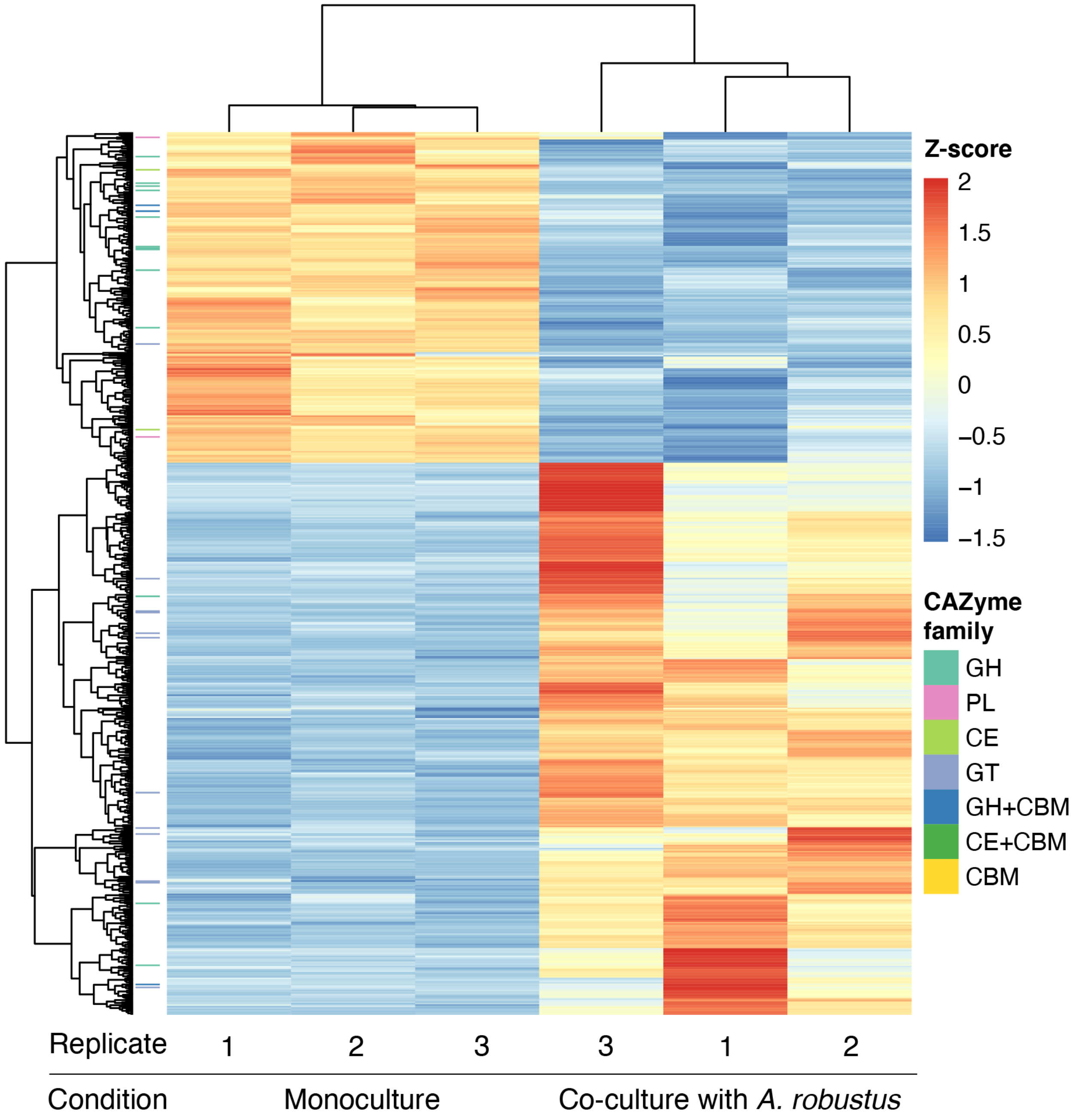


**Supplementary Figure S8. Heatmap of all differentially expressed genes for *F. succinogenes* strain UWB7 grown in monoculture *vs.* in co-culture with *A. robustus*.** Heatmap shows all the differentially expressed bacterial genes for three independent replicates when *F. succinogenes* strain UWB7 is grown in monoculture *vs.* in co-culture with *A. robustus*. Genes are colored on the left side of the heatmap based on the CAZyme family they are associated with.

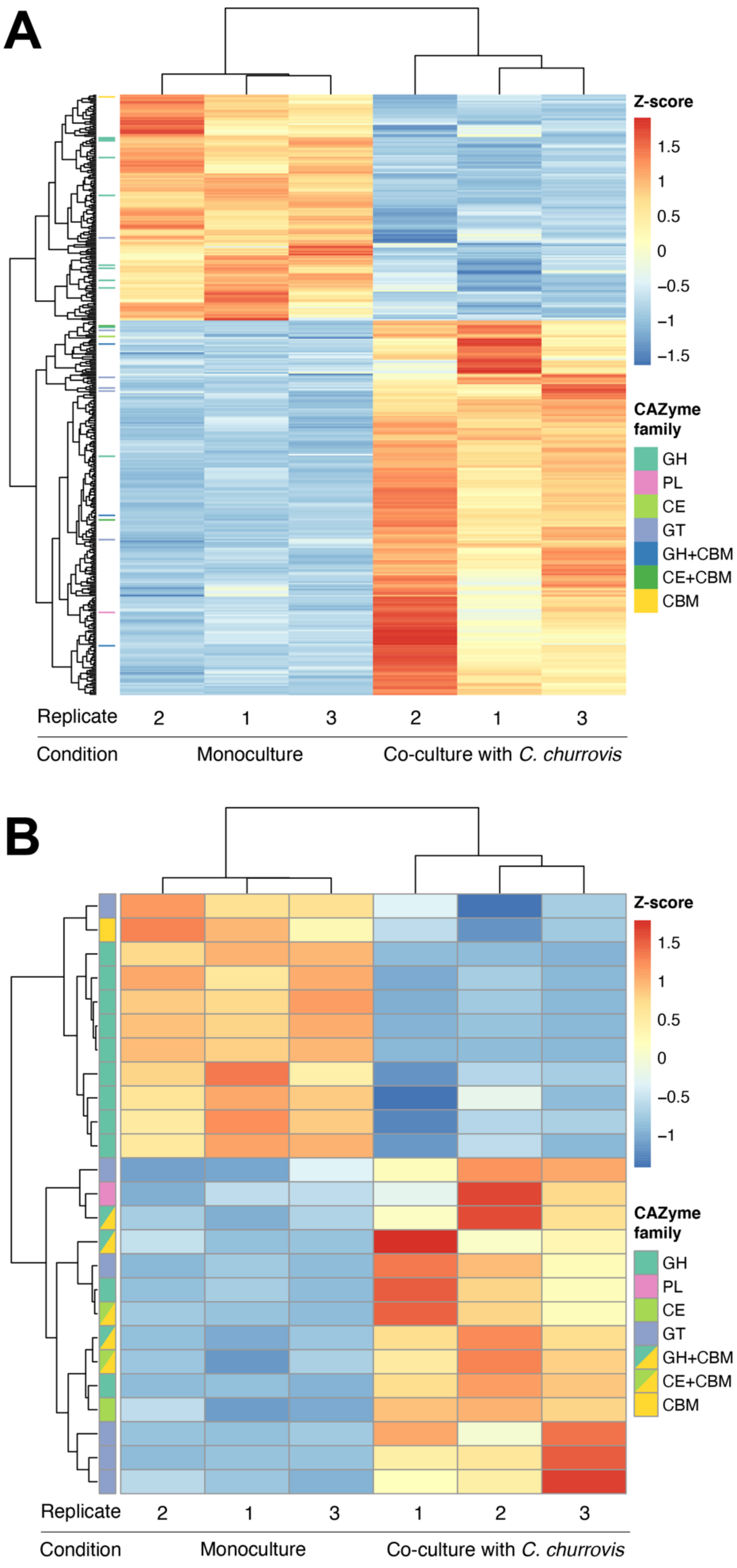
A
Z-score
1.5
1
0.5
0
−0.5
−1
−1.5
CAZyme family
GH
PL
CE
GT
GH+CBM
CE+CBM
CBM
Replicate
2
1
3
2
1
3
Condition
Monoculture
Co-culture with C. churrovis
B
Z-score
1.5
1
0.5
0
−0.5
−1
CAZyme family
GH
PL
CE
GT
GH+CBM
CE+CBM
CBM
Replicate
2
1
3
1
2
3
Condition
Monoculture
Co-culture with C. churrovis

**Supplementary Figure S9. Heatmaps of differentially expressed genes when *F. succinogenes* strain UWB7 is grown in monoculture *vs.* in co-culture with *C. churrovis*.** (**A**) Heatmap showing all differentially regulated genes for experiments performed in triplicate. (**B**) Heatmap showing only differentially regulated CAZymes for experiments performed in triplicate. Genes are colored on the left side of the heatmap based on the CAZyme family they are associated with.

**Supplementary Table S1.** Comparison between various rRNA depletion approaches.

| Supplier or publication | Working principle | Cost/sample (approx.) | Number of primers per species | Percent rRNA left | Notes |
|---|---|---|---|---|---|
| This work | Blocked reverse transcription & RNase H digestion | $10 | 20 | 1-10% (monocultures), 20% (co-cultures) | |
| Kraus *et al*, 2019 (2) | Oligo-based rRNA pull-down | N/A | 12 | <5% | |
| Culviner *et al*, 2020 (3) | Oligo-based rRNA pull-down | $10 | 21 | ~20-25% | 17 of the probes were optimized to target 8 species simultaneously |
| Huang *et al*, 2020 (4) | RNase H digestion | $12.94 | 90 | <5% to ~25% | |
| Engelhardt *et al*, 2020 (5) | RNase H digestion | N/A | 90 | <5% | Targets rRNA as well as transfer-messenger RNA (tmRNA) |
| Choe *et al*, 2021 (6) | RNase H digestion | $10 | 60-130+ | <10% | |
| Prezza *et al*, 2020 (7) | Cas9-mediated cleavage of rRNA-derived cDNA | $3-7 | 120 | ~10-50% | |
| Ribo-Zero Plus Microbiome rRNA Depletion Kit (Illumina) | RNase H digestion | N/A; similar RiboZero product is $47 | / | ~20-30% | Targets common gut bacteria, including those in ATCC MSA-2002, MSA-2005, and MSA-2006. |
| RiboMinus™ Bacteria 2.0 Transcriptome Isolation Kit (Thermo Fisher) | Oligo bead-based pull down | $82 | >150 | <10% | Targets 76 bacterial species |
| rRNA Depletion Kit for Bacteria (NEBNext) | RNase H digestion | $33 | / | ~1-5% for several species, but varies. | Compatible with ~60 bacterial species with varying efficiency (NEB & User-reported data) |

**Supplementary Table S2.** List of all primers used in EMBR-seq+, including primers for library preparation associated with CEL-seq2, EMBR-seq blocking primers, and RNase H probes. Numbers in the names of the hotspot primers and probes indicate the transcript coordinate of the rRNA hotspot. This supplementary table is provided as a separate excel file.

**Supplementary Table S3.** Pearson's *r* correlation values between RPM counts for RNA-seq libraries prepared with different methods and between replicates. This supplementary table is provided as a separate excel file.

**Supplementary Table S4.** Results of differential gene expression analysis for comparisons between *F. succinogenes* strain UWB7 grown in monoculture *vs.* co-culture, using "No depletion", "EMBR-seq", "RNase H", and "EMBR-seq+" libraries. For EMBR-seq+ results, predicted CAZyme annotations from dbCAN2 are also included (columns U-Z where applicable). In the table, columns I-N (named "M1"…"C3") show RPM for each gene (calculated as raw counts * $10^6$ / sum of raw counts). Column R (named "Alpha") indicates genes that are differentially expressed between the "No Depletion" and "EMBR-seq+" libraries, and are filtered out in column S (named "Beta") In column S, a value of 0 indicates a gene that is not differentially expressed, -1 indicates a gene that is upregulated in co-culture, and 1 indicates a gene that is upregulated in monoculture. This supplementary table is provided as a separate excel file.

**Supplementary Table S5.** Select bacterial stress response genes upregulated in co-culture with anerobic fungi *A. robustus* (Table S5A) or *C. churrovis* (Table S5B). Individual groups of genes of interest were identified by looking at COG annotations, KEGG orthology annotations, and the CAZyme gene cluster finder (CGC). In the table, columns H-M (named "M1"…"C3") show RPM for each gene (calculated as raw counts * $10^6$ / sum of raw counts). This supplementary table is provided as a separate excel file.

**Supplementary Table S6.** Comparison of bacterial transporters upregulated in co-culture in this work and those reported in Table 2 of Swift *et al.* (1). Columns A-C contain *F. succinogenes* strain UWB7 genes reported in Table 2 of Swift *et al*. as upregulated in co-culture with *C. churrovis*. Columns D-L contain $\log_2$(Fold Change) and Padj values from our datasets (*F. succinogenes* strain UWB7 co-culture with *C. churrovis* (columns D-G) and *A. robustus* (I-L)) for the genes of interest. For our datasets, a threshold of $P_{adj} < 0.1$ and $|\log_2(Fold\ Change)| > 0.5$ was used to determine if a gene was differentially expressed. This supplementary table is provided as a separate excel file.

**Supplementary Table S7.** Upregulated CAZyme genes in monocultures of *F. succinogenes* strain UWB7 *vs*. co-cultures of *F. succinogenes* strain UWB7 with *A. robustus* or *F. succinogenes* strain UWB7 with *C. churrovis*. Monocultures and co-cultures were both grown on switchgrass.

| | **Monoculture (*F. succinogenes* strain UWB7)** | | | **Co-culture (*F. succinogenes* strain UWB7 + *A. robustus*)** | |
|---|---|---|---|---|---|
| GH | GH5 | Ga0136279_0422 | GH | GH8 | Ga0136279_1446 |
| | GH5 | Ga0136279_2901 | | GH10 | Ga0136279_3005 |
| | GH8 | Ga0136279_1768 | | GH16 | Ga0136279_0928 |
| | GH9 | Ga0136279_2267 | | GH5+CBM4 | Ga0136279_1456 |
| | GH9 | Ga0136279_2903 | GT | GT2 | Ga0136279_1712 |
| | GH10 | Ga0136279_1559 | | GT2 | Ga0136279_1713 |
| | GH11 | Ga0136279_0449 | | GT2 | Ga0136279_1715 |
| | GH23 | Ga0136279_0889 | | GT2 | Ga0136279_0012 |
| | GH26 | Ga0136279_2010 | | GT2 | Ga0136279_0879 |
| | GH74+GH74 | Ga0136279_2365 | | GT2 | Ga0136279_0852 |
| | GH10+CBM6 | Ga0136279_2195 | | GT4 | Ga0136279_0554 |
| | GH26+CBM35 | Ga0136279_2970 | | GT4 | Ga0136279_0639 |
| PL | PL1 | Ga0136279_2902 | | GT8 | Ga0136279_0876 |
| | PL1 | Ga0136279_0674 | | GT32 | Ga0136279_0638 |
| CE | CE11 | Ga0136279_1153 | | GT35 | Ga0136279_0585 |
| | CE15 | Ga0136279_2918 | | | |
| GT | GT2 | Ga0136279_1342 | | | |

| | **Monoculture (*F. succinogenes* strain UWB7)** | | | **Co-culture (*F. succinogenes* strain UWB7 + *C. churrovis*)** | |
|---|---|---|---|---|---|
| GH | GH5 | Ga0136279_0422 | GH | GH8 | Ga0136279_1446 |
| | GH5 | Ga0136279_2901 | | GH16 | Ga0136279_0928 |
| | GH8 | Ga0136279_1768 | | GH5+CBM4 | Ga0136279_1456 |
| | GH8 | Ga0136279_1318 | | GH30+CBM6 | Ga0136279_2169 |
| | GH9 | Ga0136279_2267 | | GH43+CBM6 | Ga0136279_2166 |
| | GH9 | Ga0136279_2903 | PL | PL1 | Ga0136279_0688 |
| | GH10 | Ga0136279_1559 | CE | CE6 | Ga0136279_1612 |
| | GH11 | Ga0136279_0449 | | CE6+CBM6 | Ga0136279_2170 |
| | GH26 | Ga0136279_2010 | | CE6+CBM6 | Ga0136279_2171 |
| GT | GT2 | Ga0136279_1342 | GT | GT2 | Ga0136279_1712 |
| CBM | CBM51 | Ga0136279_2699 | | GT2 | Ga0136279_1713 |
| | | | | GT8 | Ga0136279_0876 |
| | | | | GT32 | Ga0136279_0638 |
| | | | | GT35 | Ga0136279_0585 |

**REFERNCES**


1. Swift CL, Louie KB, Bowen BP, Hooker CA, Solomon KV, Singan V, Daum C, Pennacchio CP, Barry K, Shutthanandan V, Evans JE, Grigoriev IV, Northen TR, O'Malley MA. 2021. Cocultivation of anaerobic fungi with rumen bacteria establishes an antagonistic Relationship. mBio 12:e01442-21.
2. Kraus AJ, Brink BG, Siegel TN. 2019. Efficient and specific oligo-based depletion of rRNA. Sci Rep 9:12281.
3. Culviner PH, Guegler CK, Laub MT. 2020. A simple, cost-effective, and robust method for rRNA depletion in RNA-sequencing studies. mBio 11:e00010-20.
4. Huang Y, Sheth RU, Kaufman A, Wang HH. 2020. Scalable and cost-effective ribonuclease-based rRNA depletion for transcriptomics. Nucleic Acids Res 48:e20.
5. Engelhardt F, Tomasch J, Häussler S. 2020. Organism-specific depletion of highly abundant RNA species from bacterial total RNA. Access Microbiol 2:acmi000159.
6. Choe D, Szubin R, Poudel S, Sastry A, Song Y, Lee Y, Cho S, Palsson B, Cho B-K. 2021. RiboRid: A low cost, advanced, and ultra-efficient method to remove ribosomal RNA for bacterial transcriptomics. PLoS Genet 17:e1009821.
7. Prezza G, Heckel T, Dietrich S, Homberger C, Westermann AJ, Vogel J. 2020. Improved bacterial RNA-seq by Cas9-based depletion of ribosomal RNA reads. RNA 26:1069–1078.
